# Enabling Model-based Design for closed-loop applications in neuroengineering

**DOI:** 10.1101/2023.09.11.557151

**Authors:** M. Di Florio, Y. Bornat, M. Caré, V. R. Cota, S. Buccelli, M. Chiappalone

## Abstract

This study addresses the inherent difficulties in the creation of neuroengineering devices for closed-loop stimulation, a task typically characterized by intricate and technically demanding processes. Beneath the substantial hardware advancements in neurotechnology, there is often rather complex low-level code that poses challenges in terms of development, documentation, and long-term maintenance. To overcome these obstacles, we adopted an alternative strategy centered on Model-based Design (MBD) as a means to simplify the creation of closed-loop systems and reduce the entry barriers. MBD offers distinct advantages by streamlining the development workflow and facilitating the implementation of intricate systems. In this study, we applied MBD techniques to implement a spike detection algorithm on a Field-Programmable Gate Array (FPGA) using commercially available hardware that combines neural probe electronics with programmable FPGA-based hardware. The entire process of data handling and data processing was designed within the Simulink® environment, with subsequent generation of HDL code tailored to the FPGA hardware. The validation of our approach was conducted through in vivo experiments involving six animals. We have made all project code files open-source, thereby providing free access to fellow scientists interested in the development of closed-loop systems.

## Introduction

Due to significant investments made by research institutions and medical companies in advancing neurotechnologies for brain disorder treatments [1], contemporary data acquisition techniques now enable the simultaneous recording from thousands of electrodes [2]. Notably, recently developed invasive recording systems offer exceptional spatial and temporal resolution, even down to the single-cell level [3], [4], thus expanding the horizons of in-vivo research. These technological enhancements have spurred substantial advancements in the fields of bioelectronics and neural engineering. Consequently, they have paved the way for the creation of cutting-edge Brain Computer Interfaces (referred to as BCI for simplicity), neuroprosthetic devices, and innovative ‘electroceutical’ approaches [5]. Electroceutical is a recently defined application field, which consists of treatments based on ‘electrical’ rather than ‘pharmacological’ therapy. Electrical therapies have sparked interests among researchers, industries and funding agents [6], [7]. Electroceutical approaches have already been successfully employed to neurological diseases, such as depression, epilepsy, gastroparesis and arrhythmia, as reported in the literature [8], [9].

Electroceutical therapy can be administered in two ways: by means of open-loop stimulation (i.e., OLS) and closed-loop stimulation (i.e., CLS). OLS implies the delivery of a stimulation pattern without considering the brain’s current state, whereas CLS allows to modulate the electrical stimuli based on the brain’s ongoing activity [10]. Indeed, closed-loop based-systems were proven to be highly advantageous in several aspects, including biological safety and therapeutic efficacy. With the goal of restoring motor function following stroke or traumatic brain injury, Guggenmos and colleagues [11] used a closed-loop approach inspired by Hebbian plasticity mechanisms. In their novel system, electrical stimuli were applied to the primary somatosensory area (S1) in a time-locked synchronous one-to-one manner to the firing of neurons on the rostral forelimb area (RFA; the pre-motor area equivalent in rats). Termed ADS (Activity Dependent Stimulation), the approach has been successfully used to improve the performance of rats in reaching and grasping tasks by functionally reconnecting the sensorimotor loop.

The exploitation of closed-loop paradigms has led to an increase in demand for high-performance signal processing techniques for hardware implementation for both clinical practice and basic research applications. It also implies the development of complex real-time algorithms to cope with the higher number of recording sites needed to study the complex spatial and temporal dynamics of the nervous system [12], [13].

The realization of closed-loop systems is a demanding process which involves three main steps: (i) recording neural activity at various levels of temporal and spatial resolution and risk; (ii) decoding or interpreting the neural activity using real-time signal processing algorithms; (iii) generating an electrical pattern of stimuli based on the output of the processing technique. Field-Programmable Gate Arrays (FPGAs) are often the first choice to create highly reconfigurable systems for different purposes while maintaining very high performance. However, the complexity involved in designing an FPGA architecture for closed-loop implementations (e.g. reading, processing, and writing the neural code [14]) can be daunting, especially for not expert scientists. To this end, it is necessary to provide simpler and faster ways to perform FPGA-based processing. Indeed, lowering the entry barrier for developing closed-loop systems is crucial to meet the growing demand and accelerate research progress.

The primary objective of this work is to address the existing technological gap by introducing a rapid-prototyping Model-Based Design architecture for accelerating the implementation of closed-loop systems on FPGAs. The focus is on testing this architecture on a commercially available system for in vivo closed-loop stimulation experiments. To enhance the capabilities of the system, a spike detection algorithm has been developed thanks to the custom modification of the original code. This change aimed at improving the system’s capability by accurately detecting spikes in real-time. The testing phase of the project was crucial in evaluating the performance and effectiveness of the architecture in real-time conditions. By comparing the offline and online implementations of the same spike detection algorithm, we proved the system’s ability to accurately detect spikes in real-time.

This work represents the first demonstration of how a neuroengineered system, developed through a Model-Based Design approach, can have a transformative impact in decreasing the developmental time. By leveraging on the advantages of this design approach, such as rapid prototyping and efficient optimization, this paper showcases the potential of model-based design methodologies in advancing the field of neuroengineering.

The full project, including modified Verilog code, Simulink® architecture, library, and documentation, is available on GitHub at the following link: https://github.com/MattiaDif/closed-loop-neuroengineering.

## Methods

### 1. General philosophy of the approach

A Model-Based Design approach, utilizing the Simulink® environment in conjunction with the HDL coder, could be a game-changer in the neuroscientific and neuroengineering research. Simulink® is a well-established tool for Model-Based Design provided by MathWorks that enables simulation of system behavior and troubleshooting of models before hardware implementation. Furthermore, the HDL code can be automatically generated from the model, speeding up the implementation process. This architecture aims to streamline the development process and enhance the efficiency of implementing closed-loop systems on FPGA platforms. The work has achieved this goal through the following steps (see Fig. 1 Panel A):

**Fig. 1.**
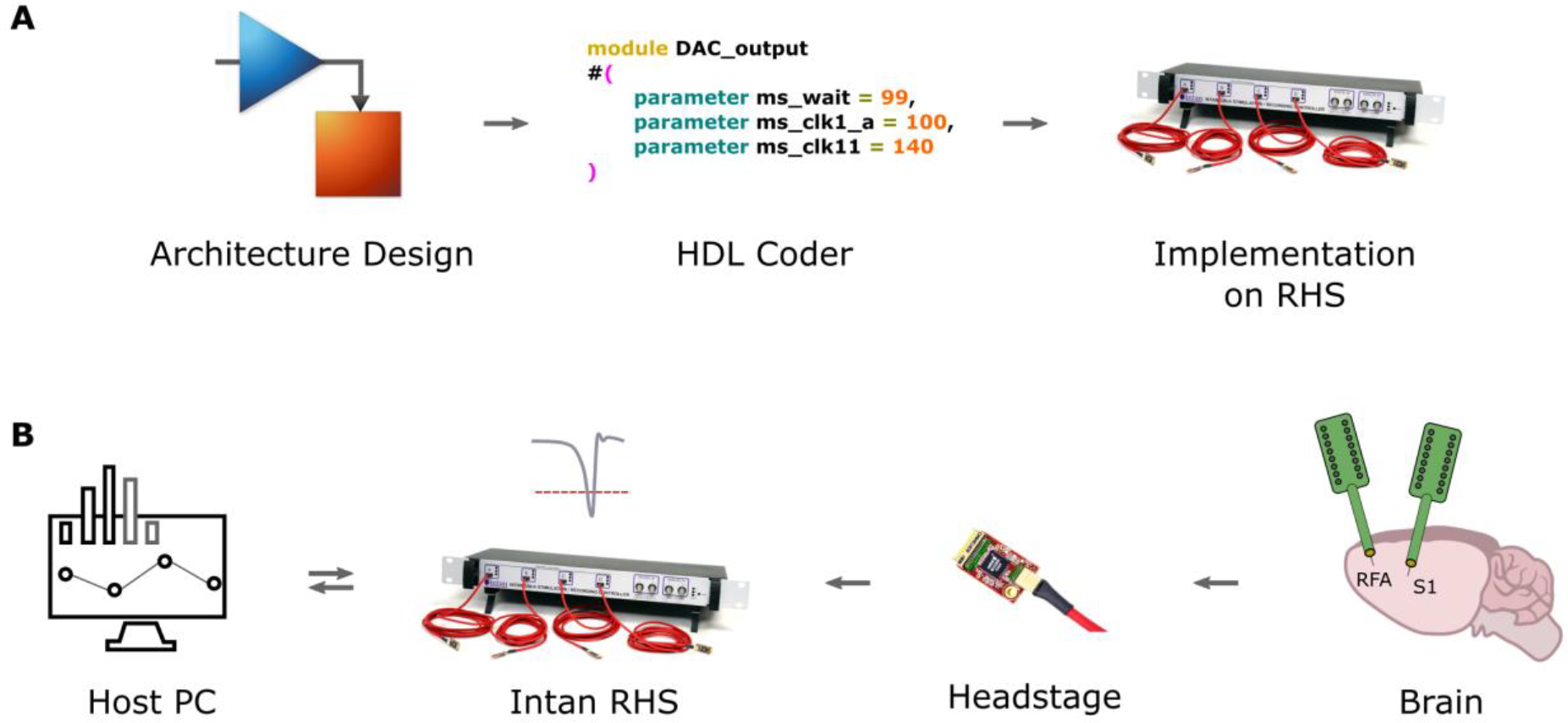
System overview and design steps. A: main steps followed for the Model-Based Design approach. All the architecture has been designed in Simulink, then, exploiting the HDL Coder, Verilog code has been generated and integrated into the original Verilog architecture of the Intan RHS. B: system used for real-time spike detection. A host PC serves as running a Graphical User Interface for online data visualization and control of the Intan RHS parameters. The Intan RHS consists of the core of the system. It is an FPGA-based controller capable of real-time data acquisition and stimulation. By modifying the original code of the controller, a spike detection algorithm has been implemented. The headstage serves as interface between the controller and the electrodes implanted in the animal brain. It amplifies the signal and provides analog to digital conversion.

1. Creating a Simulink® model for processing neural data. Simulink®, being a graphical programming environment, allows for the modeling and simulation of dynamic systems in a faster and more intuitive way with respect to traditional methodologies. By creating a comprehensive Simulink® model, the project establishes a foundation for implementing closed-loop systems on FPGAs;
2. Using HDL Coder to implement the model on FPGA: HDL Coder is employed to generate synthesizable Verilog and VHDL code from the Simulink model which can then be programmed onto the FPGA;
3. Establishing use case for commercially available systems. This use case provides real-world examples of how the architecture can be leveraged in various domains, highlighting its versatility and potential benefits. The Intan Stimulation/Recording system has been selected as the appropriate platform for the study.

### 2. System overview

The neural signal processing system adopted for the work is the Intan RHS Stim/Recording System. It is an FPGA-based commercially available data acquisition and stimulation system designed for in vivo experiments (see Fig. 1 Panel B). It consists of three main components:

1. Headstage: the headstage incorporates the Intan Technologies RHS2116 microchip. It serves as a bidirectional electrophysiology interface system and facilitates the connection of the microelectrode array. The headstage comprises 16 stimulation/amplifier blocks, which are responsible for amplifying and processing neural signals.
2. Acquisition System: the acquisition system is built around the Intan Technologies RHS Stim/Recording controller. This system is FPGA-based and designed specifically for electrophysiology data acquisition. It enables the simultaneous recording of incoming neural signals from the headstage and facilitates communication of stimulation parameters through the interface.
3. Host PC: the host PC serves as a general-purpose computer that runs software for real-time data visualization and facilitates custom parameter communication. It acts as the central control unit for the overall system. The software running on the host PC provides the necessary tools and interfaces for visualizing the recorded neural data and enables communication with the acquisition system for configuring and controlling stimulation parameters. The host PC acts as an essential interface between the user and the real-time neural signal processing system.

These three components form a full system for real-time neural data acquisition and stimulation. The headstage captures neural signals from the microelectrode array, the acquisition system handles the recording and stimulation parameters, and the host PC provides a user-friendly interface for controlling the FPGA architecture and visualizing the data in real-time.

### 3. Identifying the best Spike Detection Algorithm

The objective was to evaluate and identify the most suitable algorithm to be implemented on the FPGA. It is important to note that this step did not involve the HDL code generation directly. Instead, its primary focus was on algorithm evaluation and selection for the target embedded implementation, as reported in our previous work [15]. Based on the results of the performance analysis and considering the remaining resources available in the FPGA, a meticulous evaluation of spike detection performance and implementation complexity led to the selection of a hard threshold algorithm. This algorithm effectively identifies local peaks in the signal, striking a balance between spike detection accuracy, implementation simplicity and chip area usage.

### 4. Architecture conceptualization and design

#### 4.1 Intan RHS Verilog architecture customization

The Custom Architecture (CA) is a modified version of the original Verilog code (see Fig 2 Panel A1) provided by the Intan Technologies © company, designed to hold the Verilog code generated by the Simulink model. The CA redirects acquired data to a custom path to process the neural data in real-time. By incorporating a switch in the original path, the dataflow can be diverted through the custom path that houses the custom Verilog code generated from Simulink® (see Fig 2 Panel A1).

**Fig. 2.**
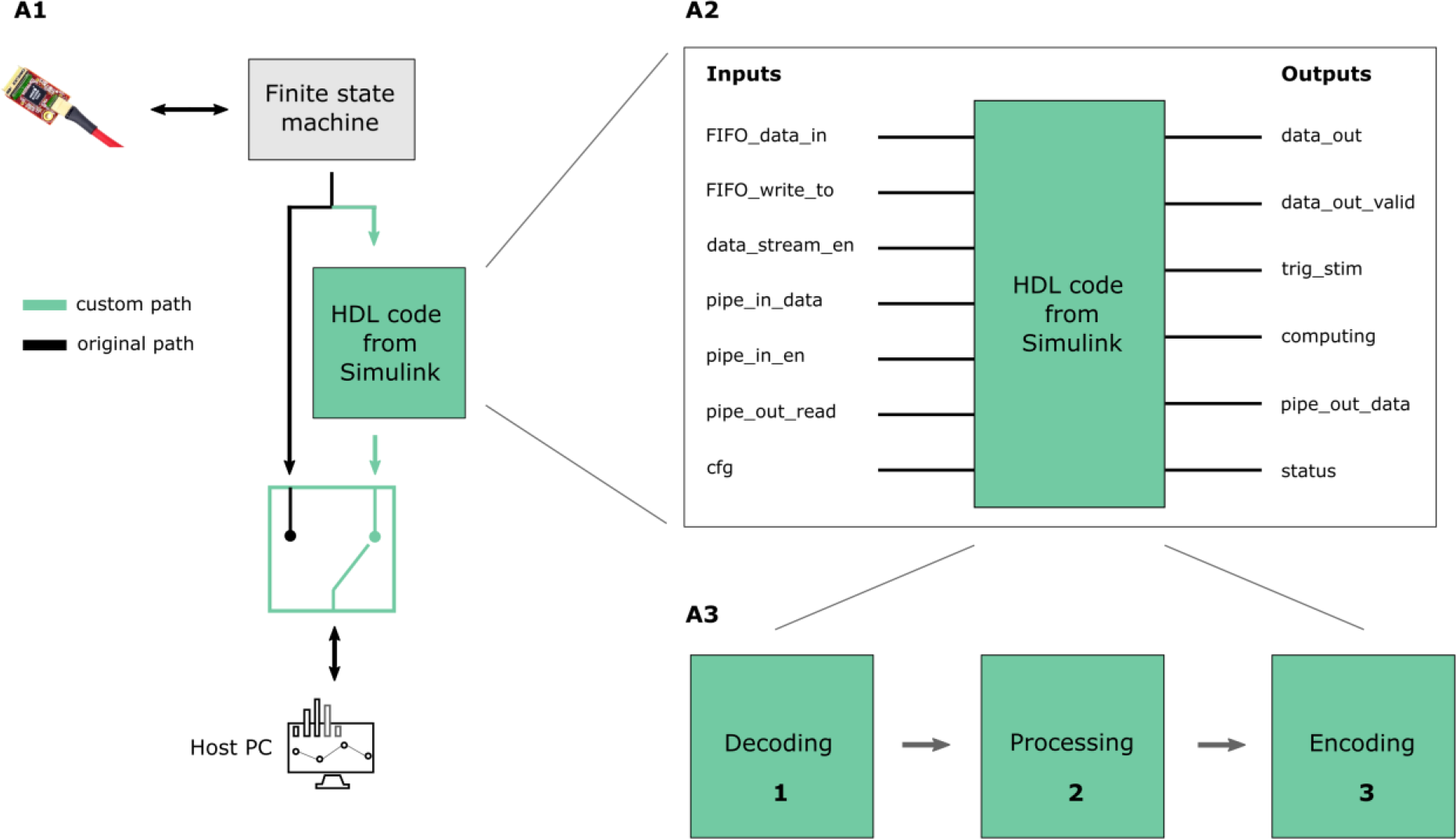
Custom architecture overview. A1: modification of the original Verilog code of the Intan RHS to redirect the dataflow through the custom path without affecting the initial behavior. The “HDL code from Simulink” block consists in the Verilog code generated from the Simulink model. A2: detail of the I/O interface of the Simulink model. This series of inputs and outputs allow communication between the output of the finite state machine and the custom paths. A3: the HDL block provides custom computation which consists of decoding, processing (filtering and spike detection) and encoding steps.

To better understand the CA architecture, it’s important to have a brief overview of the original architecture of the Intan RHS system.

The Intan RHS Stim/Recording controller is an open-source commercial device that enables recording and stimulation from 128 electrodes at the same time. The system’s core is the Opal Kelly XEM6010-LX45, a USB 2.0 integration module based on the Xilinx Spartan-6 FPGA. Opal Kelly provides a FrontPanel SDK for configuring communication and interfacing the FPGA with a PC, Mac, or Linux hardware, reducing the effort required to build functional prototypes.

Intan Technologies offers a host computer application programming interface (API), written in C++, for multi-platform support (Qt), which handles communication with the FPGA and displays the graphical user interface (GUI). Additionally, they provide hardware description language (HDL) code, written in Verilog, that manages real-time communication with the RHS2116 digital electrophysiology stimulation/amplifier chips and the host computer. The RhythmStim code, developed for the Opal Kelly board, enables streaming of up to 128 amplifier channels from multiple RHS2116 chips, data from up to 8 other ADCs, and signals from up to 16 digital inputs. It allows for setting stimulation protocols for all 128 channels, with settings such as amplitude, pulse width, and sequences configurable through the GUI. The data is synchronized, time stamped, and transmitted over a standard USB 2.0 bus to the host computer at a rate exceeding 20 MByte/s. The core of the RhythmStim code is a Finite State Machine (FSM) that cycles through 140 repeating states to execute a single serial peripheral interface (SPI) cycle. Each SPI cycle involves executing a command like “convert(0)” to retrieve the actual value of channel “0” of all the connected headstages, and transferring the data to the FIFO within the same 140 states. A counter is then incremented from 0 to 19 to track the series of 20 commands sent to the RHS2116 (16 “convert” commands for each amplifier channel and 4 auxiliary commands for enabling stimulation channels and other operations). This FSM is the principal component of the overall original architecture, allowing to handle the sampled data from the headstage redirecting them to the Host PC. By intercepting this stream of data, we were able to process the samples in real-time.

#### 4.2 Inputs/Outputs interface

To guarantee communication between the original and custom paths, an input/output interface has been designed using Verilog (see Fig 2 Panel A2).

List of input ports:

- FIFO_data_in: this input represents the recorded data from the headstage;
- FIFO_write_to: this input is a flag that indicates the validity of the retrieved data. If the value is 1, the data retrieved is informative. Otherwise, it is 0;
- data_stream_en: this is an 8-bit register that provides information about the enabled headstages and the port they are connected to on the controller. A value of 1 in position 0 indicates that the first headstage is connected. A value of 1 in position 0 and in position 1 indicates that the first and second headstages are connected and will send values;
- pipe_in_data: this input contains the pipe-in endpoint values that come from the Qt GUI. This port is useful for sending custom parameters from the host PC;
- pipe_in_en: this input contains the pipe-in enable endpoints that come from the Qt GUI. If the value is 1, it means that a command was sent through pipe-in data. This port informs the FPGA about the validity of the data sent from the host PC;
- pipe_out_read: this input is currently unused and left for future development;
- cfg: this input is left for future development and is intended to inform the custom architecture about the current configuration version.

List of output ports:

- comp_data_out: this port contains the recorded data to be transmitted to the Host computer via the USB interface;
- comp_data_out_valid: this boolean port becomes true when comp_data_out contains valid data to be transmitted via the USB interface;
- trig_stim: this port is reserved for future development and will send triggers for stimulation to close the loop;
- computing: this port is designed to inform external modules that the custom architecture is still computing and not yet ready to process other tasks. It is currently not in use;
- pipe_out_data: this endpoint is intended for debugging. It is useful for sending informative data from the FPGA to the Host PC;
- status: this port will indicate the status of the custom architecture module.

In summary, the signal dataflow is redirected through the custom path where real-time processing takes place. This custom path is built using Verilog code generated by a Simulink model, allowing users to customize the Verilog architecture and implement their own algorithms. At present, the custom processing involves a high-pass filter and hard threshold spike detection which looks for a local peak in the signal.

#### 4.3 Custom signal processing

The signal processing algorithm was developed using the Simulink® HDL library provided by MathWorks. After the model design, Verilog code was generated and integrated into the modified Verilog architecture. The input/output interface acts as a communication bridge between the Verilog code and the Simulink model.

The custom computation developed in Simulink consists of three primary steps: data decoding, data processing, and data encoding (see Fig. 2 Panel A3):

1. Data Decoding: This initial step involves the interpretation and conversion of input data into a format that can be readily processed by the subsequent stages. If the input data is in a specific encoding or compressed format, it needs to be decoded and transformed into its original representation. In the custom computation developed in Simulink, the input data is first decoded using a Moore State Machine implemented with StateFlow. StateFlow is a graphical language within Simulink that encompasses various tools such as state transition diagrams, flow charts, state transition tables, and truth tables. The purpose of the Moore State Machine is to determine the channel and headstage from which the input data was acquired. It operates by transitioning between different states based on the input data and current state. The output of a Moore machine is solely determined by the current state it is in. During each time step, the Moore chart computes its output, evaluates the input data, and configures itself for the next time step accordingly. By utilizing the StateFlow implementation, the input data is decoded and the relevant information regarding the channel and headstage from which the data originated is extracted. This approach provides a structured and organized manner to handle the decoding process, enabling the system to accurately determine the source of the input data.
2. Data Processing: Once the input data is decoded, it undergoes various processing operations to extract relevant information and perform the desired computations. This step typically involves applying algorithms, filters, transformations, or any other necessary operations to manipulate and analyze the data according to the intended purpose. Within the data processing block of the architecture, there are two main steps: filtering the raw data and performing spike detection computations. The first step involves filtering the raw data that is sampled by the headstage. In the current implementation, a 2nd order highpass Butterworth filter is utilized. This filter is designed to remove low-frequency components from the signal, allowing for the preservation of higher-frequency components. The cutoff frequency of the filter is set at 300 Hz. Once the data is filtered, spike detection is performed using a hard threshold spike detection algorithm. In this algorithm, a local peak, specifically a negative one, is identified as an action potential (see Figure 3). The spike detection is based on a comparison of the current sample at time t, the sample at time t-1, and the sample at time t-2. To identify a spike, two conditions must be met: (i) the sample at time t-1 must be the lowest among the three samples, and (ii) the sample at time t-1 must be lower than a set threshold. If both conditions are satisfied, a spike is identified. Once a spike is identified, a refractory period begins. The refractory period is a specific duration of time during which the system ignores any potential action potentials that occur. This period is implemented to ensure that subsequent spikes are not erroneously detected in close temporal proximity to the initial spike. Additionally, it is worth noting that the architecture of the processing block is fully serial, meaning that the filtering and spike detection operations are performed on one channel at a time. This design choice was made to effectively manage the available resources within the FPGA. By processing one channel at a time, the architecture optimizes resource utilization. Although the testing was conducted on 32 channels, the architecture has the capacity to process up to 128 channels in real-time. The maximum sampling frequency achievable is 30 kHz, specifically 20kHz, 25kHz or 30kHz can be selected in accordance with the original architecture.
3. Data Encoding: after the processing stage, data encoding is employed to reconstruct the output data flow, ensuring that the transmitted data is in the appropriate format to be received and interpreted correctly by the Host PC GUI. This encoding step is crucial for maintaining the integrity and structure of the processed data during transmission. To enable the storage of filtered data and timestamps of the identified spikes, the dataflow to the host PC was customized. While this customization provides the advantage of retaining this valuable information about the filtered data and spike timestamps, it came at the expense of the raw data.

**Fig. 3.**
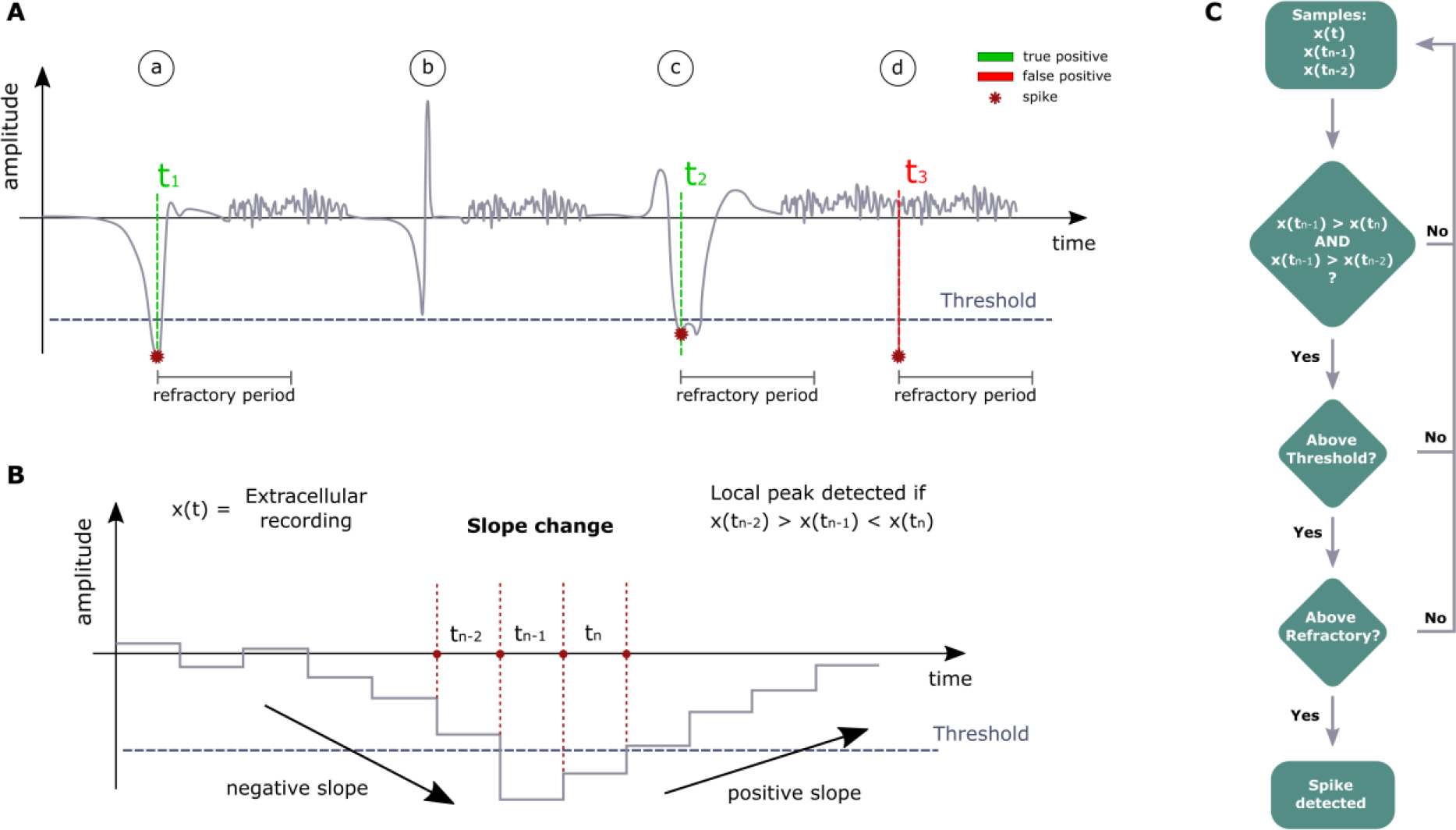
Spike detection algorithm functioning schematic. A: examples of true positive, false negative and false positive detections of the algorithm. a) an action potential below the threshold is identified correctly (true positive). b) an action potential above threshold is not identified (false negative). c) an action potential which shows a double peak is correctly identified just once because of the refractory period that begins as soon as the first peak is detected, avoiding a double detection (true positive). d) a noisy peak below threshold is wrongly detected by the algorithm (false positive). B: details of algorithm functioning. For the peak detection, the current sample at time t_n_, the previous sample at time t_n-1_ and the sample at time t_n-2_ are considered. In the sample at time t_n-1_ is below threshold and it is the lowest, a spike is identified. C: flow chart to summarize the main steps of the algorithm.

**Fig. 4.**
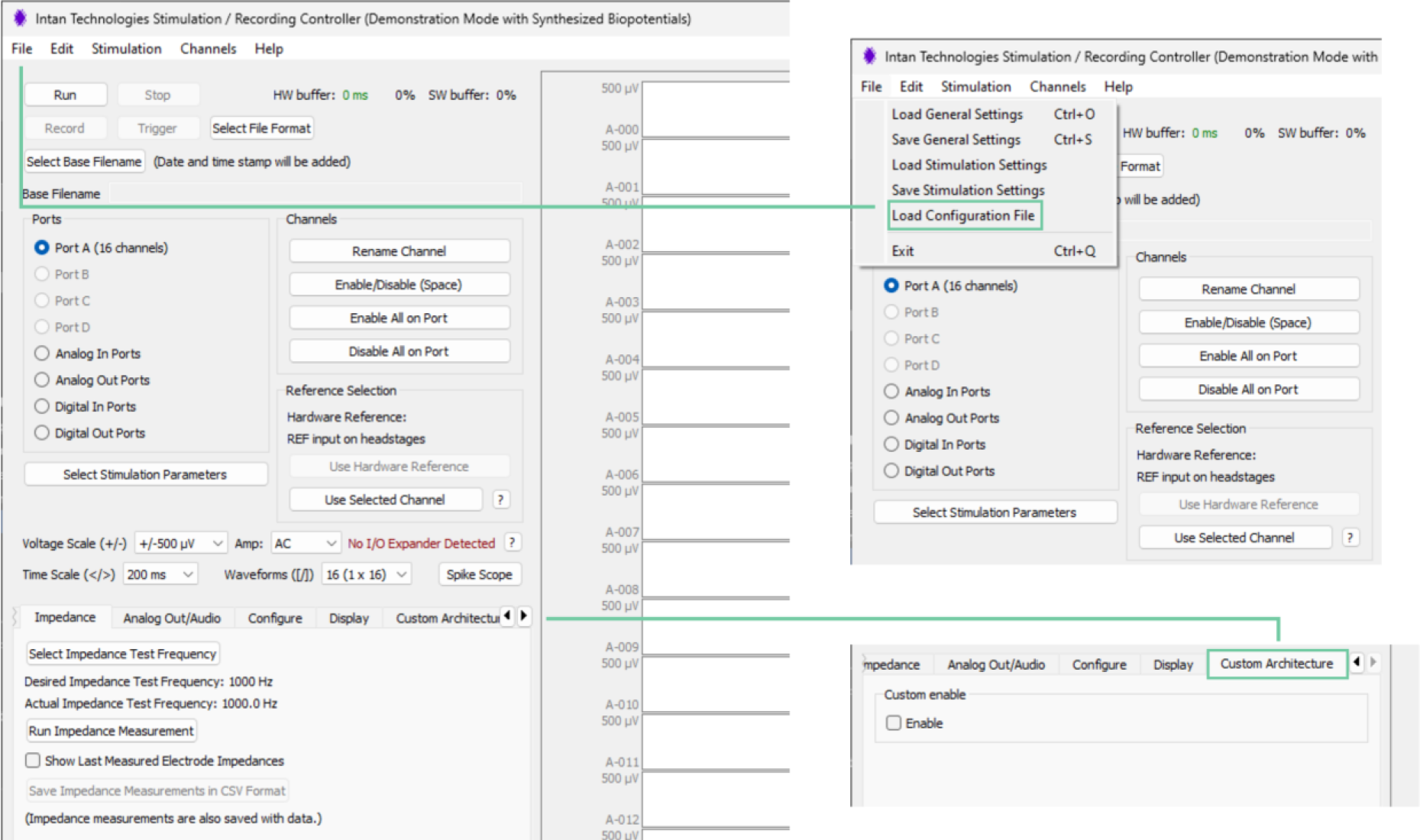
Customized GUI. An item called “Load Configuration File” in the File windows has been added to allow for the loading of a configuration file which includes the threshold value and the refractory period to apply. A custom tab called “Custom Architecture” has been added with a flag to enable the dataflow redirection to the custom path. If this flag is not enabled, the system maintains its original behavior.

**Fig. 5.**
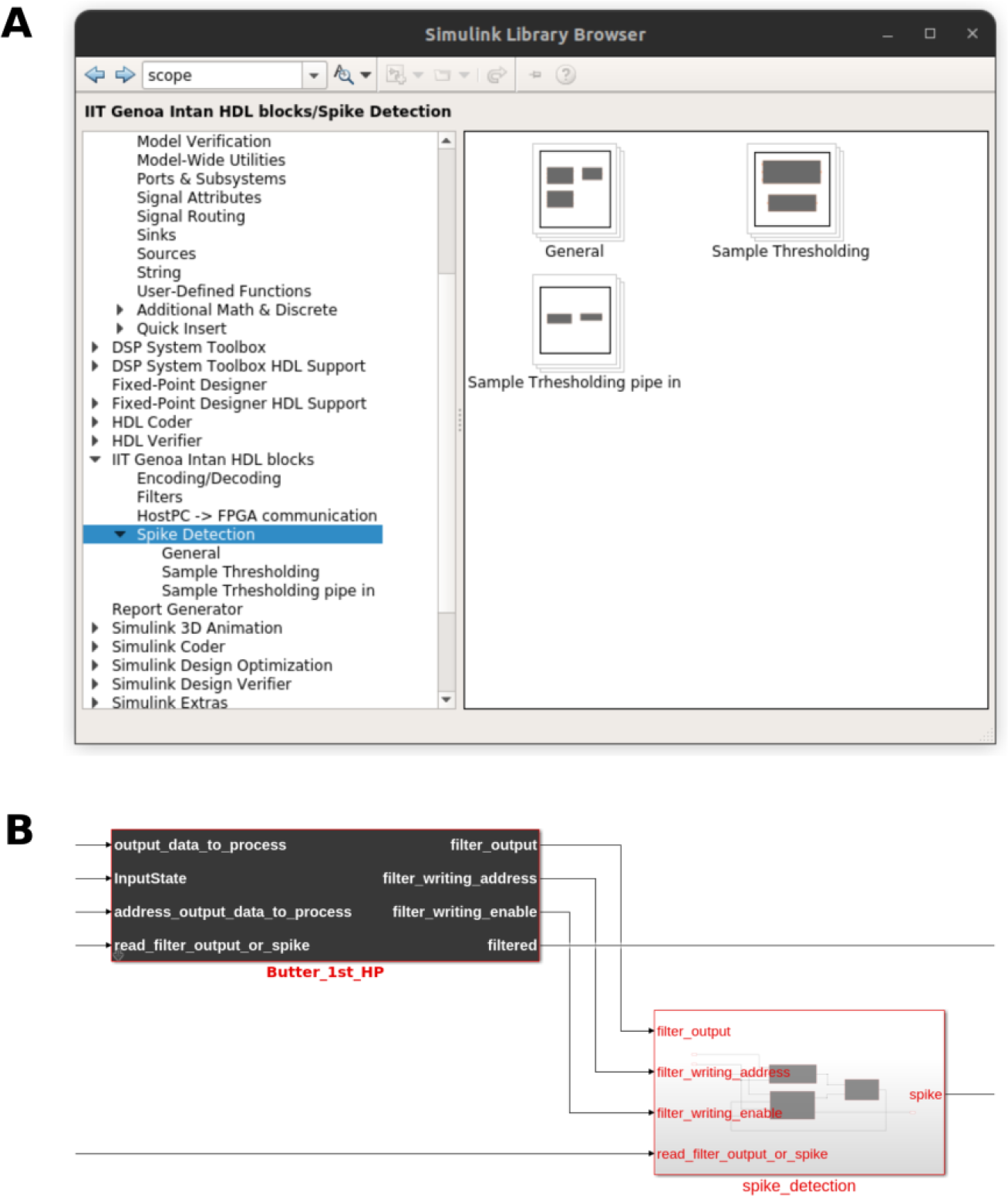
Simulink® library. A: library developed for Simulink®. It can be installed once downloaded the GitHub repository. It provides specific block for the design of spike detection algorithms already tested and validated for the Intan RHS. B: Example of application. A user can drag and drop its desired blocks into the Simulink® environment and, following the name of the I/O ports, he can intuitively connect them

By following this sequence of data decoding, processing, and encoding, the custom computation in Simulink can effectively transform the input data into a processed output that is appropriately formatted, ready to be transmitted to the host PC. It is important to highlight that each block, especially the signal processing block, has been designed to be as independent as possible from one another. This design approach allows for easy replacement of these blocks with user-defined blocks, providing flexibility and customization options. By designing the blocks to be modular and independent, the project enables users to replace or modify specific components according to their specific needs and requirements.

### 5. Graphical user interface customization

To enable the rerouting of the dataflow through the custom path, and the delivery of custom parameters to the FPGA, the manufacturer’s original open-source C/C++ code has been modified (see Figure 6). These modifications allow for the integration of the custom computation within the existing codebase. The original graphical user interface (GUI), which was developed using Qt 5.8, serves as the front-end for the USB interface of the Opal Kelly XEM6010-LX45. The GUI has been extended to include a new tab called “Custom_arch.” Within this tab, there is a flag named “Cst_en” that can be toggled to activate the custom computation. When the “Cst_en” flag is set, a specific register of the architecture is written, triggering the switch to the custom architecture. This mechanism ensures that the dataflow is redirected through the custom computation path when the flag is enabled (see Fig. 2 Panel A1). In the File windows of the system’s GUI, a new item called “Load Configuration File” has been added. This functionality allows users to specify custom threshold values and refractory periods for the spike detection computation. By selecting this option, users can load a text file organized in a specific format. The text file contains the customized threshold and refractory period values for the spike detection algorithm. It is structured in a way that facilitates easy loading of the custom parameters. However, at the current state of the project, the same threshold and refractory period values are applied to all channels. This feature provides flexibility for users to adjust the spike detection parameters according to their experimental conditions.

**Fig. 6.**
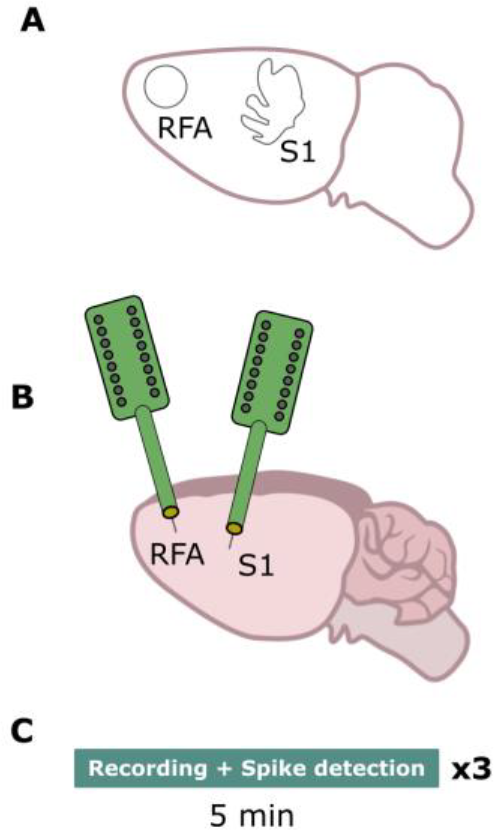
Experimental protocol. A: identification of the areas of interest by means of a stereotaxic frame. B: placement of the electrode arrays in the rostral forelimb area (RFA) and in the primary somatosensory area (S1). C: timeline of experiments includes 5 minutes recordings and real-time spike detection from both areas per 3 times. Each time a different threshold has been used based on the experimental conditions (i.e. background noise level and spikes amplitude).

The software, including the modified GUI and the custom computation code, has been compiled using Qt Creator community edition. It has been specifically compiled for the Windows 10 64-bit operating system, ensuring compatibility with the target environment. These modifications and adaptations enable the integration of the custom computation within the existing software infrastructure, allowing for the seamless execution and control of the custom architecture through the modified GUI.

### 6. Simulink® library

To make closed-loop system development more accessible to non-experts, a custom Simulink library has been developed (see Fig. 5). This library contains preconfigured blocks that are specifically designed to interface with the Intan RHS system, simplifying the development process. The custom Simulink library includes a collection of blocks that have been preconfigured with the necessary parameters and settings to seamlessly integrate with the Intan RHS system. These blocks can be easily connected and configured within the Simulink environment, allowing users to design and customize their own algorithms without requiring specialized knowledge in FPGA programming or hardware design.

### 7 Experiments

#### 7.1 Animals

The protocol for animal use was approved by the Italian Ministry of Health and Animal Care (authorization n. 509/2020-PR). The study included acute experiments conducted on 6 healthy adult Long-Evans rats (male, weighing between 300-400g, aged 4-5 months) from Charles River Laboratories, Calco, LC. Upon performing neuroscientific experiments for another study [16], we then recorded spontaneous neural activity before the conclusion of the session. The experimental procedures were conducted at the Animal Facility of the Italian Institute of Technology (IIT), Genova, Italy, and were approved by the Italian Ministry of Health and Animal Care (authorization n. 509/2020-PR).

#### 7.2 Surgical procedures

Rats were anesthetized by placing them inside a vaporizing chamber and injecting gaseous isoflurane (5% @ 1 lpm). To achieve surgical level anesthesia, ketamine (80-100 mg/kg IP) and xylazine (5-10 mg/kg IM) were administered. The rat was then secured in a stereotaxic frame, and vital parameters were monitored throughout the procedure. A midline skin incision was made after applying lidocaine cream as a topical analgesic. A laminectomy was performed successfully at the level of the Cisterna Magna to allow the cerebrospinal fluid (CSF) to drain. Burr holes (3 mm diameter) were made over the primary somatosensory area (S1) and rostral forelimb area (RFA) based on stereotaxic measurements [9] at –1.25, +4.25 and +3.5, +2.5 AP, ML, respectively. Finally, the dura mater was removed from both burr holes to allow insertion of MEAs (Neural Probes; A4x4-5 mm-100-125-703-A16, NeuroNexus).

#### 7.3 Experimental protocol and recording paradigm

The placement of the MEAs was carried out following the analysis of the blood vasculature pattern and based on stereotaxic coordinates. A 16-channel probe (NeuroNexus A4x4-5mm-100-703-A16) was implanted in both RFA and S1. The experimental protocol involved recording basal activity for 5 minutes from both electrodes. Real-time filtering and spike detection were performed enabling the CA (see Figure 2). The extracellular signals were continuously sampled at a depth of 16 bits and a rate of 20 kHz or 30 kHz. For the spike detection algorithm, a refractory period of 1.5ms was set and different thresholds ranging from -50 μV to -110 μV were configured based on the noise level of the background noise and the spikes amplitude. For each animal 3 different thresholds [th1, th2, th3] have been set equal to [-50 μV, -70 μV, - 90 μV] or [-70 μV, -90 μV, -110 μV].

### 8 Offline performance testing

An exact replica of the hard threshold spike detection algorithm has been developed in MATLAB for offline comparison of the logic. The performance of spike detection was evaluated by comparing the results of offline spike detection with real-time spike detection using in vivo data. The objective was to ensure that the spike detection results obtained in real-time were identical to the results obtained from the offline analysis. This comparison involved examining the number and positions of the detected spikes with both implementations.

## Results

A thorough comparison between offline and real-time implementation of the spike detection algorithms was conducted for each of the 32 channels across the 6 animals in the study. The offline hard threshold algorithm was applied to the dataset and the results compared to the output of the real-time implementation as shown in Fig. 7.

**Fig. 7.**
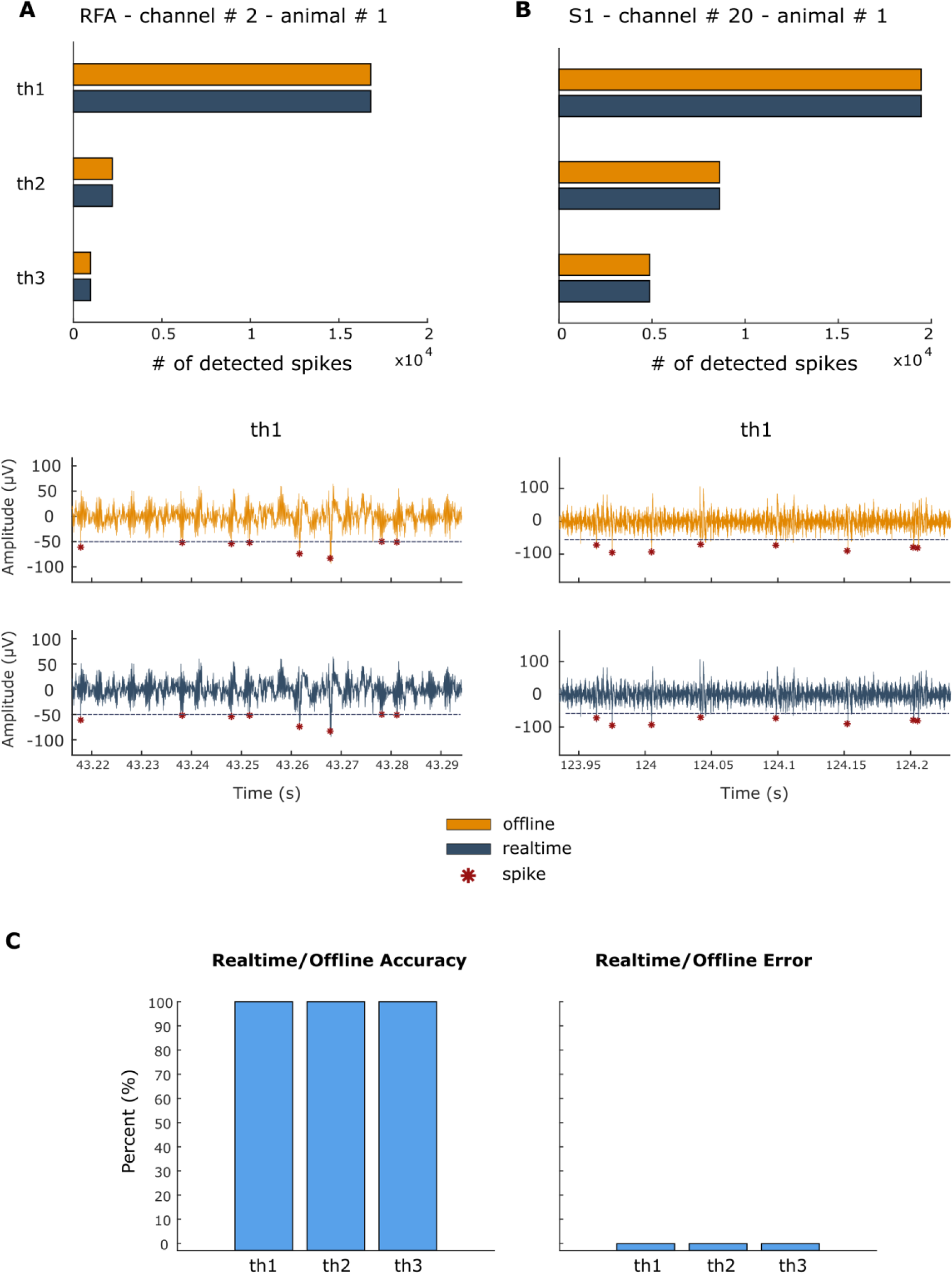
Panel A-B: results for channel # 2 and channel # 20 of animal # 1. Panel C: results overview. The orange color is referred to the offline output, while the blue color to the real-time one. A: number of detected spikes in RFA area for the 3 thresholds used (th1 = -50μV, th2 = -70μV and th3 = -90μV). The number of spikes is always identical between offline and real-time implementation. Below, a zoom of the spike detected for the two implementations for th1. B: number of detected spikes in S1 area for the 3 thresholds used (th1 = -50μV, th2 = -70μV and th3 = -90μV). The number of spikes is always identical between offline and real-time implementation. Below, a zoom of the spike detected for the two implementations for th1. C: These values have been obtained by averaging the accuracy and error values for each channel of each animal. On the left, the accuracy of the real-time over the offline is reported. A 100% accuracy is obtained in each case, showing that the real-time implementation behaves as the offline algorithm. On the right, the percentage error is shown. As acknowledged by the accuracy percentage, a 0% error has been obtained in each case.

In Fig. 7 Panel A and B, exemplifying plots demonstrate the offline behavior versus the real-time behavior. For both areas, RFA and S1, have been compared the number of spikes identified by the implementation. In each case, the exact number of spikes and the exact same spikes in the signal are detected for the different thresholds.

To quantify the performance of the real-time spike detection, two metrics have been calculated: i) percentage accuracy and ii) percentage error. The percentage accuracy is defined as:

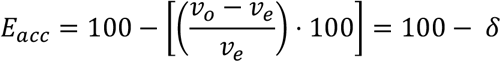

where *E*_*acc*_ is the accuracy (or accuracy error), *v*_*o*_ is the observed value (total number of spikes detected by the real-time algorithm), *v*_*e*_ is the expected value (total number of spikes of offline algorithm) and δ is the percentage error. The percentage error is defined as:

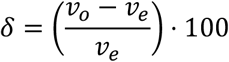

where δ is the percentage error, *v*_*o*_ is the observed value and *v*_*e*_ is the expected value.

The accuracy and error have been computed for each channel of each of the 6 animals and the overall average has been computed. As shown in Fig. 7 Panel C, we achieved 100% accuracy, then 0% error.

## Discussion

One of the key advancements achieved through this work is the enhancement of the system’s capabilities in terms of filtering and spike detection in neural signals. By leveraging the Model-Based Design methodology, the system was able to achieve accurate and reliable spike detection in real-time, identical to its offline implementation. This improvement in performance demonstrates the effectiveness of the design workflow in enhancing the system’s functionality. The successful implementation of the Model-Based Design approach in this project serves as a compelling example of how this methodology can be leveraged for the design and development of advanced neurotechnologies in general.

The project adopted a strategic approach that aimed to leverage an existing commercial system for neurophysiology, rather than designing a system from scratch. By utilizing the capabilities of the commercial system, the focus was primarily on the development of custom computation modules to enhance its functionality [17]. The commercial system, in this case, was the Intan Stimulation/Recording system, which provided a solid foundation for neurophysiological data acquisition and stimulation.

The Model-Based Design approach enables the development of complex architectures with a clear and structured design flow. In contrast to conventional methods [18]–[20], our work has demonstrated several notable advantages, building upon the experience gained during its execution. The approach we employed offers numerous benefits, including a high level of abstraction, ease of implementation, and the ability to intuitively validate and verify designs through comprehensive simulation and testing procedures.

This modular and independent design philosophy enhances the versatility and adaptability of the overall architecture. It empowers users to tailor the signal processing pipeline to their specific application or research objectives, without the need for extensive modifications or reconfigurations of the entire system. Furthermore, this design approach promotes code reusability and facilitates collaboration within the research community. Researchers can easily share and exchange custom-designed blocks, promoting the advancement and refinement of signal processing techniques in the field of closed-loop neuroscience.

The decision to prioritize the storage of filtered data and spike timestamps over raw data preservation is a trade-off that depends on the specific requirements and objectives of the project. Indeed, while having access to the filtered data and spike timestamps can be valuable for various analysis and post-processing purposes, the inability to retrieve the raw data means that certain types of analysis or further investigations relying on the unprocessed data may not be possible. It is possible to modify the front-end architecture to allow the storage of raw data alongside the filtered data and timestamps of the detected spikes. By incorporating appropriate data storage mechanisms and memory resources, the raw data can be preserved for further analysis or reference.

At the current state of the project, the hard threshold algorithm used for spike detection could be considered a simplistic approach. However, it is widely used at the core of many spike sorting algorithm [21]. There is certainly room for further improvement by implementing more advanced spike detection algorithms [22]. However, careful consideration should be given to the computational complexity, memory requirements, and real-time processing constraints imposed by the FPGA platform. The algorithm’s efficiency and compatibility with the available resources should be evaluated to ensure seamless integration.

The validation process for the architecture involved testing it on a subset of 32 channels out of the total 128 available channels. The reason for this limitation was the lack of sufficient chips during the experiments. Based on the results obtained from the offline testbenches, it is evident that the architecture is capable of correctly processing and detecting spikes in neural signals when operating with up to 128 channels.

It is worth mentioning that although the current implementation applies identical threshold and refractory period values across all channels, future enhancements and updates to the project may allow for individual channel customization if desired.

The application of Model-Based Design in the neurotechnology field shows great potential based on the outputs and achievements of this project. By leveraging a model-based approach, the development and implementation of complex architectures for neurotechnology systems have been streamlined, enabling faster and more efficient design processes. Furthermore, it has facilitated the validation and verification of the developed architecture. Through testbenches and comparisons with offline analysis, the accuracy and reliability of the real-time implementation have been confirmed. The potential of this methodology in the neurotechnology field lies in its ability to accelerate the development cycle, thus enabling more advanced and sophisticated systems. By providing a unified modeling and more intuitive platform, it promotes collaboration and knowledge sharing among researchers, lowering the entry barrier in the design and development of such systems.

In face of the above-described achievement, it is our understanding that the technology described in this work represents a significant contribution to the development of closed-loop neurotechnologies.

This project is actively being developed and is available to the community. As part of this ongoing development, new features and enhancements will be introduced over time to expand and improve the current capabilities. By making the project available to the community, it not only encourages collaboration and knowledge sharing but also opens up opportunities for contributions from the wider user base. This community-driven approach will help drive innovation and enable the system to evolve in line with the needs and requirements of its users.

## Acknowledgment

The authors wish to express their gratitude to all the colleagues who directly or indirectly contributed to the making of this project. A special thanks to the colleagues Vijay Iyer and Akshay Rajhans from MathWorks, who mentored the work. V.R.C. was supported by Marie Skłodowska-Curie Individual Fellowship MoRPHEUS, Grant Agreement no. 101032054, funded by the European Union under the framework programme H2020-EU.1.3.-EXCELLENT SCIENCE.

This work received financial support from MathWorks and was carried out within the framework of the project “RAISE - Robotics and AI for Socio-economic Empowerment” and has been supported by European Union – NextGenerationEU.

## Bibliography

[1] M. Chiappalone et al., “Neuromorphic-Based Neuroprostheses for Brain Rewiring: State-of-the-Art and Perspectives in Neuroengineering,” Brain Sci., vol. 12, no. 11, p. 1578, Nov. 2022, doi: 10.3390/brainsci12111578.

[2] E. T. Zhao et al., “A CMOS-based highly scalable flexible neural electrode interface,” Sci. Adv., vol. 9, no. 23, p. eadf9524, 2023, doi: 10.1126/sciadv.adf9524.

[3] N. A. Steinmetz et al., “Neuropixels 2.0: A miniaturized high-density probe for stable, long-term brain recordings,” Science (80-.)., vol. 372, no. 6539, pp. 139–148, 2021, doi: 10.1126/science.abf4588.

[4] G. N. Angotzi et al., “SiNAPS: An implantable active pixel sensor CMOS-probe for simultaneous large-scale neural recordings,” Biosens. Bioelectron., vol. 126, no. August 2018, pp. 355–364, 2019, doi: 10.1016/j.bios.2018.10.032.

[5] M. Semprini et al., “Technological approaches for neurorehabilitation: From robotic devices to brain stimulation and beyond,” Front. Neurol., vol. 9, no. APR, pp. 1–9, 2018, doi: 10.3389/fneur.2018.00212.

[6] K. Famm, B. Litt, K. J. Tracey, E. S. Boyden, and M. Slaoui, “A jump-start for electroceuticals,” Nature, vol. 496, no. 7444, pp. 159–161, Apr. 2013, doi: 10.1038/496159a.

[7] S. Reardon, “Electroceuticals spark interest.,” Nature, vol. 511, no. 7507, p. 18, 2014, doi: 10.1038/511018a.

[8] G. Panuccio, M. Semprini, L. Natale, S. Buccelli, I. Colombi, and M. Chiappalone, “Progress in Neuroengineering for brain repair: New challenges and open issues,” Brain Neurosci. Adv., vol. 2, p. 239821281877647, 2018, doi: 10.1177/2398212818776475.

[9] T. Denison, M. Morris, and F. Sun, “Smart neural stimulators sense and respond to the body’s fluctuations,” IEEE Spectr., 2015.

[10] T. Levi, P. Bonifazi, P. Massobrio, and M. Chiappalone, “Editorial: Closed-loop systems for nextgeneration neuroprostheses,” Front. Neurosci., vol. 12, no. FEB, pp. 10–12, 2018, doi: 10.3389/fnins.2018.00026.

[11] D. J. Guggenmos et al., “Restoration of function after brain damage using a neural prosthesis,” Proc. Natl. Acad. Sci. U. S. A., vol. 110, no. 52, pp. 21177–21182, 2013, doi: 10.1073/pnas.1316885110.

[12] Y. Tchoe et al., “Human brain mapping with multithousand-channel PtNRGrids resolves spatiotemporal dynamics,” Sci. Transl. Med., vol. 14, no. 628, 2022, doi: 10.1126/scitranslmed.abj1441.

[13] C. Zrenner, D. Desideri, P. Belardinelli, and U. Ziemann, “Real-time EEG-defined excitability states determine efficacy of TMS-induced plasticity in human motor cortex,” Brain Stimul., vol. 11, no. 2, pp. 374–389, 2018, doi: 10.1016/j.brs.2017.11.016.

[14] A. Mazzoni and M. Chiappalone, “Editorial: Reading and writing the neural code for neuroprosthetics,” Front. Neurosci., vol. 17, 2023, doi: 10.3389/fnins.2023.1130921.

[15] M. Di Florio, V. Iyer, A. Rajhans, S. Buccelli, and M. Chiappalone, “Model-based online implementation of spike detection algorithms for neuroengineering applications,” Proc. Annu. Int. Conf. IEEE Eng. Med. Biol. Soc. EMBS, vol. 2022-July, pp. 736–739, 2022, doi: 10.1109/EMBC48229.2022.9871444.

[16] M. Carè, F. Barban, M. Di Florio, R. Greco, C. Tassorelli, and M. Chiappalone, “Towards personalized electroceutical therapy: electrophysiological investigations in a pre-clinical model of ischemic lesion,” Proc. GNB 2023, pp. 1–4, 2023.

[17] M. Murphy et al., “Improving an open-source commercial system to reliably perform activity-dependent stimulation,” J. Neural Eng., vol. 16, no. 6, 2019, doi: 10.1088/1741-2552/ab3319.

[18] M. Tambaro, M. Bisio, M. Maschietto, A. Leparulo, and S. Vassanelli, “FPGA Design Integration of a 32-Microelectrodes Low-Latency Spike Detector in a Commercial System for Intracortical Recordings,” Digital, vol. 1, no. 1, pp. 34–53, 2021, doi: 10.3390/digital1010003.

[19] S. Gibson, J. W. Judy, and D. Marković, “An FPGA-based platform for accelerated offline spike sorting,” J. Neurosci. Methods, vol. 215, no. 1, pp. 1–11, 2013, doi: 10.1016/j.jneumeth.2013.01.026.

[20] E. A. Vallicelli et al., “Neural spike digital detector on FPGA,” Electron., vol. 7, no. 12, 2018, doi: 10.3390/electronics7120392.

[21] H. G. Rey, C. Pedreira, and R. Quian Quiroga, “Past, present and future of spike sorting techniques,” Brain Res. Bull., vol. 119, pp. 106–117, 2015, doi: 10.1016/j.brainresbull.2015.04.007.

[22] F. Lieb, H. G. Stark, and C. Thielemann, “A stationary wavelet transform and a time-frequency based spike detection algorithm for extracellular recorded data,” J. Neural Eng., vol. 14, no. 3, p. aa654b, 2017, doi: 10.1088/1741-2552/aa654b.

